# Impact of two neighbouring ribosomal protein clusters on biogenesis factor binding and assembly of yeast late small ribosomal subunit precursors

**DOI:** 10.1101/398149

**Authors:** Jan Linnemann, Gisela Pöll, Steffen Jakob, Sébastien Ferreira-Cerca, Joachim Griesenbeck, Herbert Tschochner, Philipp Milkereit

## Abstract

Many of the small ribosomal subunit proteins are required for the stabilisation of late small ribosomal subunit (SSU) precursors and for final SSU rRNA processing in *S. cerevisiae*. Among them are ribosomal proteins (r-proteins) which form a protein cluster around rpS0 (uS2) at the “neck” of the SSU (S0-cluster) and others forming a nearby protein cluster around rpS3 (uS3) at the SSU “beak”. Here we applied semi-quantitative proteomics together with complementary biochemical approaches to study how incomplete assembly of these two r-protein clusters affects binding and release of SSU maturation factors and assembly of other r-proteins in late SSU precursors in *S. cerevisiae*. For each of the two clusters specific impairment of the local r-protein assembly state was observed. Besides, cluster-specific effects on the association of biogenesis factors were detected.

These suggested a role of S0-cluster formation for the efficient release of the two nuclear export factors Rrp12 and Slx9 from SSU precursors and for the correct incorporation of the late acting biogenesis factor Rio2. Based on our and on previous results we propose the existence of at least two different r-protein assembly checkpoints during late SSU maturation in *S. cerevisiae*. We discuss in the light of recent SSU precursor structure models how r-protein assembly states might be sensed by biogenesis factors at the S0-cluster checkpoint.

## Introduction

The small ribosomal subunit (SSU) plays an essential and universally conserved role during cellular translation of mRNA into proteins. In the unicellular eukaryote *S. cerevisiae*, hereafter called yeast, the SSU consists of 33 ribosomal proteins (r-proteins) and the 18S ribosomal RNA (rRNA). Current atomic resolution structure models suggest that the fold of the 18S rRNA backbone largely defines the conserved overall architecture of the yeast SSU [1]. Hence, the 5′ and the central 18S rRNA secondary structure domains form the SSU “body” with the “platform” while the 18S rRNA 3′ major domain forms the SSU “head”. RNA-RNA interactions in between the secondary structure domains stabilize their spatial orientation towards each other. In addition, r-proteins further define the inter-domain configuration, besides their architectural role inside the individual rRNA secondary structure domains. As an example rpS0 (uS2 according to the r-protein nomenclature published in [2]), rpS2 (uS5) and rpS21 (eS21) establish one protein cluster (hereafter called S0-cluster) at the head body junction which establishes a link between the three major secondary structure domains through interactions with helix 1 in the 5′ domain, helices 25 and 26 in the central domain and helices 28, 35 and 36 in the 3′ major domain. This connecting point is further reinforced by RNA-RNA base pairing interactions in the central pseudoknot which are formed between the loop originating from helix 1 and the region just before the 3′ major domain, to which rpS2 (uS5) directly binds. The biogenesis pathway of the SSU was intensively studied in yeast (reviewed in [3]). Here, as in other eukaryotes, it starts in the nucleolus with the transcription of rRNA genes into large precursor transcripts. These contain the SSU 18S rRNA, followed by the large ribosomal subunit (LSU) 5.8S rRNA and 25S rRNA. In these precursor transcripts the 18S rRNA region is flanked at its 5′ end by the 5′ external transcribed spacer (5′ETS) and at its 3′ end by the internal transcribed spacer 1 (ITS1) preceding the LSU 5.8S rRNA region (see S1 Fig). Early during SSU maturation nucleobase and ribose modifications are introduced in the 18S rRNA region in a highly specific manner. In addition, the 5′ETS region is removed by endonucleases and the ITS1 region is cleaved. This generates the 20S rRNA precursor (pre-rRNA) of the 18S rRNA and separates the SSU and LSU precursor particles. A large number of ribosome biogenesis factors and some small nucleolar RNAs (snoRNAs) as the U3 snoRNA are required for these early steps of rRNA processing to occur in a productive fashion. Interestingly, structural and functional studies corroborated that the 5′ ETS region is not solely by-product of 18S rRNA but rather fulfils crucial functions early during SSU maturation (reviewed in [4,5]). It first serves as platform for the initial recruitment of numerous ribosome biogenesis factors and of the U3 snoRNA. Secondly, it provides a structural scaffold for the resulting large RNP which organizes the early architecture of 18S rRNA precursors. Importantly, the ensemble of 5′ETS rRNA, U3 snoRNA and associated factors keep the nascent SSU rRNA secondary structure domains apart from each other during early SSU maturation steps. This is in part due to the action of U3 snoRNA which prevents formation of the central pseudoknot by base pairing with the respective SSU rRNA regions.

After successful generation of the 20S pre-rRNA, most of the early acting factors are released together with the U3 snoRNA as well as the 5′ ETS from the particle. Thus, at this point only a few ribosome biogenesis factors stay associated with the generated late nuclear SSU precursors. These include Enp1, Dim1, Pno1, Hrr25, Nob1, Slx9 and Rrp12 [6–12]. Together with three additional late recruited factors, Rio2, Ltv1 and Tsr1, they are thought to accompany the SSU precursors through the nuclear pore to the cytoplasm [6,13–15]. Rrp12 might facilitate nuclear export by direct interactions with the nuclear pore, Slx9 and Rio2 by recruitment of the exportin Crm1/Xpo1 [11,12,16]. Besides, all the late associating factors were shown to be essential for either nuclear or cytoplasmic SSU pre-rRNA stabilisation and processing [7,11–13,15,17–22]. Recent single molecule cryo electron microscopy (cryo-EM) analyses of yeast cytoplasmic 40S precursors revealed the mode of binding of some of these factors [23–26]. From this it appears that, besides their primary requirement for pre-rRNA stabilisation, some cytoplasmic factors might have adopted additional roles in preventing the engagement of immature cytoplasmic SSUs in translation initiation and elongation [24]. Thus, the SSU binding site of Rio2 overlaps with the binding sites of the translation initiation factor eiF1A and of the initiator tRNA with eiF2. Furthermore, the binding of Tsr1 seems incompatible with the binding of eiF1A and eiF3, while Pno1 partially occupies the SSU mRNA channel [27]. In line with this, sedimentation analyses of yeast extracts indicate that mature SSUs are more efficiently recruited into the pool of translating ribosomes than their 20S pre-rRNA containing precursors [28]. In the cytoplasm members of this group of late factors are released apparently in a stepwise and in part hierarchical manner. Briefly, factors as Rrp12 and Slx9 are thought to leave the SSU precursors shortly after nuclear export, and factors as Rio2, Tsr1, Enp1, Dim1, Hrr25 and Ltv1 at intermediate cytoplasmic stages. The latest immature SSUs still contain the factors Pno1 and Nob1 together with Rio1, which joins late in the cytoplasm [29,30]. The kinase Hrr25 was implicated in the release of Ltv1, Enp1 and Tsr1 [23,31–33], the adenylate kinase Fap7 in the release of Dim1 [34], Ltv1 in the release of Tsr1 [21], and ATP consumption by Rio2 and Rio1 was implicated in their own release and the one of several of the other factors [35–38].

Surprisingly, the *in vivo* depletion of late acting factors such as Tsr1 and Fap7 [24,34,39] as well as the interference with (late) factor removal [36] or the inhibition of the pre-18S rRNA endonuclease Nob1 [40] can all lead to formation of “80S like” particles through association of immature SSUs with mature large ribosomal subunits. In some of these situations 80S like particles are devoid of mRNA and tRNA and thus seem not to represent canonical translation intermediates [39]. By contrast, *in vivo* depletion of the presumably very lately released proteins Rio1 and Nob1 partly abolishes the exclusion of immature SSUs from the translating pool of ribosomes [28]. In any case, *in vitro* and *in vivo* evidence suggests that the final SSU rRNA cleavage at the 3′ end of 18S rRNA by the PIN domain type endonuclease Nob1 can be stimulated in one or the other way by SSU precursor interactions with mature large ribosomal subunits [40,41].

Previous work indicated that most of the yeast SSU r-proteins play important and specific roles for SSU pre-rRNA processing and transport steps and that yeast SSU precursor RNAs are prone to degradation upon shortage of supply with SSU r-proteins (reviewed in [42]). Eliminating misassembled precursors already during SSU maturation might be an economic way to increase the proportion of cytoplasmic ribosomes with accurate functional properties. One group of yeast SSU r-proteins was shown to be important for the stabilisation and processing of early nucleolar SSU pre-rRNAs. This group includes most of the binders of the 5′ and of the central domain forming the SSU body (rpS1 (eS1), rpS6 (eS6), rpS8 (eS8), rpS9 (uS4), rpS11 (uS17), rpS13 (uS15), rpS14 (uS11), rpS22 (uS8), rpS23 (uS12), rpS24 (eS24), rpS27 (eS27)), together with a few upstream binders of the 3′ major domain forming the SSU head (rpS5 (uS7), rpS16 (uS9), rpS18 (uS13), rpS19 (eS19) [42–45]. Densities for all but one (rpS19) of these SSU r-proteins could be readily observed in recent cryo-EM analyses of purified yeast early nucleolar SSU precursors [46,47]. That is consistent with a direct role of these r-proteins in early SSU maturation steps. Still, when compared to mature subunits most of them have not yet established all contacts with rRNA and/or r-proteins in these early precursors. Hence, in agreement with conclusions from many biochemical analyses their assembly seems to proceed in a stepwise fashion with progressive establishment of intra-ribosomal interactions [42]. Analysis of the protein composition of nucleolar SSU precursors produced after expression shutdown of some of these r-proteins indicated that the release of several early acting biogenesis factors from particles depleted of these r-proteins is severely delayed [44]. In addition, evidence was found that recruitment of sub-groups of early acting factors is impeded in a specific manner reflecting the SSU rRNA binding sites of the depleted r-proteins. Hence, these studies provided some mechanistic insights into how yeast SSU r-proteins affect early nuclear SSU precursor maturation and stability. While reduced recruitment of factors in particles lacking r-proteins might delay their maturation and determine them for nuclear degradation, the trapping of other factors in misassembled particles might influence the stability and maturation of newly made SSU precursors due to the shortage of these factors.

Another large group of SSU r-proteins was observed to be required for the maturation of late 20S pre-rRNA containing SSU precursors which accumulate in the cytoplasm upon their depletion (hereafter called “late acting” SSU r-proteins). One of them is rps26 (eS26), which stands out by its late assembly properties [48,49] and binds in mature ribosomes in the SSU central domain close to the final SSU pre-rRNA processing site, the 3′ end of 18SrRNA. Besides, this group of r-proteins consists of most of the SSU head domain binders (rpS3 (uS3), rpS10 (eS10), rpS15 (uS19), rpS20 (uS10), rpS28 (eS28), rpS29 (uS14), rpS31 (eS31)), including the S0-cluster proteins rpS0 (uS2), rpS2 (uS5) and rpS21 (eS21) at the head-body junction [42,43,50–52]. Many of the late acting r-proteins could not be assigned to cryo-EM based density maps of early nuclear SSU precursors and in several cases major structural remodelling seems required to allow the establishment of their final stable assembly state [46,47,49,53]. The latter is especially true for the S0-cluster. Part of its binding site is blocked in early nuclear SSU precursors by the U3 snoRNA and other factors [46,47]. Thus, formation of the central pseudoknot and of the mature spatial configuration of the three major SSU secondary structure domains at the head-body junction is prevented. Cryo-EM structure models of human SSU precursors suggest that after release of U3 snoRNA and early factors, part of the S0-cluster binding sites are then occupied by Rrp12 [54]. Apparently, these configurations are important to allow for early SSU assembly, processing and transport, while the S0-cluster r-proteins, which stabilize the corresponding mature configuration, are required for efficient later cytoplasmic SSU maturation.

We were interested to get more insights into how formation of the S0-cluster might affect late yeast SSU maturation. More precisely, we wondered whether S0-cluster formation has consequences on the recruitment or release of biogenesis factors, or on the assembly of other r-proteins. Therefore, we decided to apply a combination of proteomic and other biochemical approaches to characterize the protein composition of late SSU precursor particles in yeast conditional expression mutants of S0- cluster genes. To distinguish more global effects of incomplete assembly of late acting r-proteins from effects which are specific for S0-cluster proteins we also included another group of late acting SSU r-proteins in these analyses. These were rpS3 (uS3), rpS20 (uS10) and rpS29 (uS14), which form together with rpS10 (eS10) a protein cluster (in the following called S3-cluster) in the SSU head domain. The S3-cluster is located close to the S0-cluster at the “beak” of the head domain, and S3- cluster proteins rpS3 (uS3) and rpS20 (uS10) were recently found to play a role for the cytoplasmic release of biogenesis factors as Ltv1, Tsr1, Rio2 and Enp1 [31].

## Results

### Effects of S0-cluster formation on Rio2 association with SSU precursors

To get insights into how S0-cluster formation affects the composition of cytoplasmic SSU-precursors we made use of previously established yeast strains each conditionally expressing one of the S0- cluster proteins [43,52]. Mutants allowing for conditional expression of the nearby S3-cluster proteins rpS3, rpS20 or rpS29 were included in the analyses to distinguish local from more global effects of incomplete assembly of late acting r-proteins. In all of these strains the respective chromosomal r-protein genes are deleted and the consequent lethal phenotypes are complemented by an episomally encoded r-protein under the control of the galactose inducible GAL1/10 promoter. Hence, they can be cultivated in galactose containing medium while shift to glucose containing medium leads to the *in vivo* depletion of the particular r-protein with lethal effects on SSU maturation [43,52]. The genomes of these strains were further modified to express the SSU biogenesis factor Rio2 in fusion with a tandem affinity purification tag (TAP-tag) (see Materials and Methods). Rio2 binds in wild type condition to 20S pre-rRNA containing, mostly cytoplasmic SSU precursors [6] at a location just opposite of the S0-cluster at the subunit interface site of the head-body junction [23]. A beta hairpin in the Rio2 N-terminal winged helix turn helix (wHTH) domain is reaching underneath the SSU head towards a beta hairpin of the S0-cluster protein rpS2.

To test how S0-cluster formation affects Rio2 association with SSU precursors the corresponding yeast strains were grown in galactose containing medium and were then incubated in glucose containing medium to shut down the production of rpS0, rpS2 or rpS21. For comparison, control strains expressing tagged Rio2, and S0-cluster proteins under control of their own promoters were treated similarly. Cellular extracts were prepared and tagged Rio2 was affinity purified in buffer conditions supporting its association with SSU precursors (see Materials and Methods). Northern blotting analyses of RNA extracted from the input fractions were in agreement with the previously observed delay in processing of 20S pre-rRNA after *in vivo* depletion of S0-cluster proteins (Fig 1, compare ratio of 20S pre-rRNA to 18S rRNA in lanes 1 and 5 with the ones in lanes 2-4). As described previously, some minor effects on earlier SSU pre-rRNA processing steps were observed (Fig 1, compare amounts of 35S pre-rRNA, 33S pre-rRNA and 32S pre-rRNA in lanes 1 and 5 with the ones in lanes 2-4) [43]. As expected, RNA analyses of the affinity purified fractions showed specific association of Rio2 with 20S pre-rRNA containing SSU precursors in wild type conditions (Fig 1, lanes 6 and 10). Rio2 still co-purified these SSU precursors after expression shut down of S0-cluster proteins. However, the apparent co-purification efficiency was reproducibly reduced, especially after *in vivo* depletion of rpS2 (Fig 1, compare 20S pre-rRNA levels in lanes 6 and 10 with the ones in lane 9). In line with previous results this effect was not obvious in control experiments using strains conditionally expressing S3-cluster proteins (Fig 2, compare 20S pre-rRNA levels in lane 5 with the ones in lanes 6-8) [31,55]. We take these results as indication for S0-cluster formation and rpS2 assembly playing a specific role for the accurate integration of Rio2 into SSU precursors.

**Fig 1.**
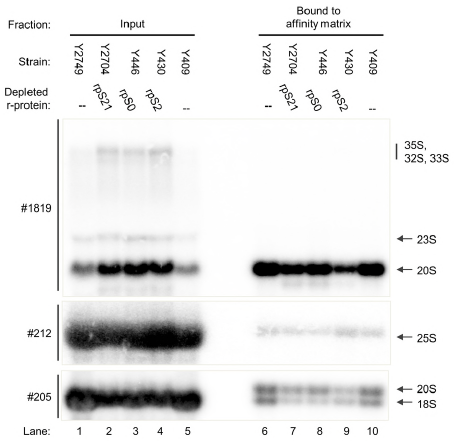
Association of Rio2 with SSU precursors depleted of S0-cluster proteins. The indicated yeast strains were cultivated for four hours in glucose containing medium to shut down expression of either rpS0, rpS2, rpS21 or no r-protein. TAP tagged Rio2 was affinity purified from cellular extracts as described in Materials and Methods. Northern blotting analyses from input and affinity purified fractions are shown (see Materials and Methods). Oligonucleotides used for RNA detection are shown on the left (#, see S1 Fig for binding sites) and detected (pre-)rRNAs are indicated on the right.

**Fig 2.**
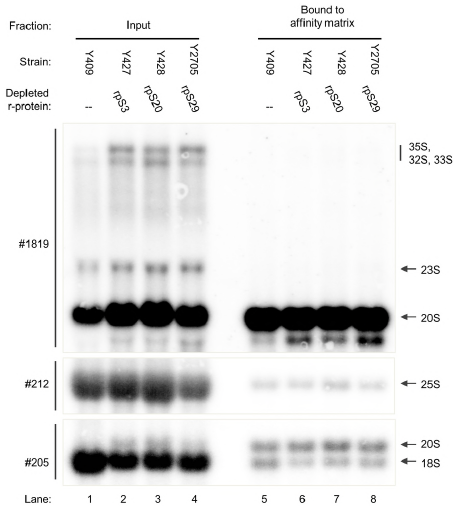
Association of Rio2 with SSU precursors depleted of S3-cluster proteins. The indicated yeast strains were cultivated for four hours in glucose containing medium to shut down expression of either rpS3, rpS20, rpS29 or no r-protein. Otherwise, see legend to Fig 1.

To further corroborate this assumption, sedimentation analyses were performed on sucrose gradients with cellular extracts prepared after expression shut down of rpS2 (Fig 3A and B). Indeed, protein analyses of the obtained fractions by western blotting indicated that the ratio between unbound Rio2 in light fractions (Fig 3B, fractions 1+2) and presumably pre-SSU associated Rio2 in heavier fractions (Fig 3B, fractions 3-6) increased when compared to wild type conditions (Fig 3B, compare right panel with left panel). RNA analyses by northern blotting suggested that 20S pre-rRNA containing SSU precursors were efficiently excluded from 80S-like particles and translating ribosomes independently of rpS2 expression (Fig 3A, left and right panel, compare 18S rRNA and 20S pre-rRNA distribution between 40S fractions 4 and 5 and 80S and polysomal fractions 6 to 11).

**Fig 3.**
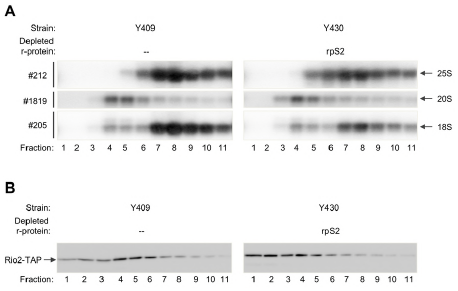
Sedimentation analyses of Rio2 and SSU precursors after *in vivo* depletion of rpS2. The indicated yeast strains were cultivated for four hours in glucose containing medium to either shut down expression of rpS2 or not. Sedimentation analyses on sucrose gradients were performed as described in Materials and Methods and part of the fractions was analysed either by northern blotting to detect (pre-)rRNAs (A) or by western blotting to detect TAP tagged Rio2 (B). In (A) oligonucleotides used for RNA detection are shown on the left (#, see S1 Fig for binding sites) and detected (pre-)rRNAs are indicated on the right.

We take these results as evidence for a role of S0-cluster formation in the proper binding of Rio2 to SSU precursors. That contrasts with SSU maturation functions of the adjacent S3-cluster, components of which seem to influence the release of Rio2 [31]. The marked impact of rpS2 among S0-cluster proteins on Rio2 binding might be linked to its beta-hairpin which closely approaches parts of the Rio2 wHTH domain.

### Changes in biogenesis factor and r-protein composition of late SSU precursors upon expression shut down of S0- or S3-cluster proteins

We were next interested to explore the possible consequences of impaired S0-cluster formation on the recruitment or release of other SSU precursor components. Therefore, we chose to undertake a more unbiased proteomic approach to compare the SSU precursor protein compositions in the respective mutant strains. *In vivo* depletion of the three S0-cluster proteins was performed as before and a control strain expressing all three proteins from their endogenous promoters was included in the experiments. Tagged Rio2 was chosen again as bait for particle purification since it enabled to enrich substantial amounts of accumulating late SSU precursors from all four strains, despite its destabilised interaction upon incomplete S0-cluster formation. Semi-quantitative mass spectrometry was used to compare pairwise the SSU precursor protein composition of the control strain with the one of each mutant strain (Materials and Methods). Similar analyses were performed with expression mutants of S3-cluster proteins rpS3, rpS20 and rpS29 to identify effects specific to one or the other r-protein cluster.

The general protein composition identified in these experiments matched well with previous protein analyses of Rio2 associated SSU precursors [6]. Most of the peptides identified in the purified fractions derived from SSU r-proteins and from SSU biogenesis factors (Figs 4A and 5A, see number of peptides in brackets). These included the bait Rio2 itself together with Pno1, Nob1, Tsr1, Ltv1, Dim1 and Enp1 which are known major components of late SSU precursors (see Introduction). For the S3-cluster expression mutants, the level of these factors remained nearly unchanged when compared to wild type conditions, with a tendency for some increase (around 25%) in case of rpS3 and rpS20 (Fig 4A, S4 File). Levels of the two factors Rrp12 and Krr1 which are believed to leave the SSU precursors early after nuclear export, significantly decreased (Fig 4A). That might indicate that their release from accumulating SSU precursors is not affected by expression shut down of S3-cluster proteins. Interestingly, also levels of the casein kinase type protein kinase Hrr25 decreased. Hrr25 was previously implicated in the release of factors as Enp1 and Ltv1 [23,31–33]. Reduced SSU precursor association of Hrr25 might interfere with this function and contribute to the observed impaired release of factors upon improper S3-cluster formation [31].

**Fig 4.**
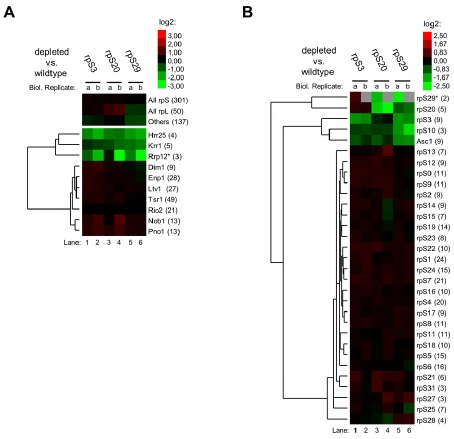
Semi-quantitative proteomic analyses of changes in the Rio2 associated SSU precursor protein composition after *in vivo* depletion of S3-cluster proteins. Yeast strains were cultivated for four hours in glucose containing medium to shut down expression of either rpS3 (Y427), rpS20 (Y428), rpS29 (Y2705) or no r-protein (Y409). SSU precursors associated with TAP tagged Rio2 were affinity purified and the SSU precursor protein composition of each conditional expression mutant was compared with the one of wildtype cells (depleted vs. wildtype) by semi-quantitative mass spectrometry as described in Materials and Methods. The results of two independent biological replicates (a, b) for ribosome biogenesis factors (A) and for SSU r-proteins (B) are visualized as heat map (colour legend in the upper right corner). Trees on the left of the heatmaps show results of hierarchical clustering analyses to identify groups of proteins with similar changes (see Materials and Methods). The respective names of proteins are indicated on the right of the heatmaps with the average number of peptides identified for each protein in brackets. A star (*) indicates that the respective protein was identified in one of the analyses just by one peptide.

Besides, the semi-quantitative proteomic analyses indicated that expression shut down of any of the tested S3-cluster proteins leads to local assembly defects insight the S3-cluster (Fig 4B). In addition, some effects on the non-essential head domain r-protein Asc1 were observed which interacts with the C-terminal extension of rpS3. RpS20 and rpS29 form interaction sites for the rpS3 N-terminal domain in the precursor SSU head domain [23]. Consistently, the apparent effects of rpS20 or rpS29 depletion on rpS3 association were stronger than the other way around. In addition, rpS10 and Enp1 binding sites in (precursor-)SSUs were previously observed to overlap [23]. Hence, effects of rps3, rpS20 or rpS29 depletion on rpS10 levels might be related to the presence of Enp1 in the accumulating SSU precursors.

The results of semi-quantitative proteomic analyses upon expression shut down of S0-cluster proteins clearly differed from those observed upon S3-cluster depletion (Fig 5, compare with Fig 4). After prolonged times of rpS2 *in vivo* depletion a pronounced relative reduction in levels of SSU r-proteins and SSU biogenesis factors was observed (Fig 5A and B, lanes 6 and 7). This further reinforced the assumption from the previous experiments that Rio2 binding to SSU precursors lacking rpS2 is weakened. Besides, after expression shut down of S0-cluster proteins the strongest reduction among all SSU r-proteins was consistently observed for the three S0 cluster r-proteins and for Asc1, rpS17 and rpS3 (Fig 5B). A clear decrease of S3 cluster proteins rpS20 and rpS29 could not be detected, but a definite statement about their assembly state remains difficult due to the comparably low peptide counts in these cases. The consistent effects inside the S0-cluster and on rpS17 were not observed in the S3-cluster mutant strains (compare Fig 5B with Fig 4B). That strongly argued for SSU assembly phenotypes differing between and being specific for the two r-protein clusters. Remarkably, the rpS17 carboxy-terminal domain extends to and intensively interacts with rpS0 at the head body junction. Hence, as before, most of the observed major effects of S0-cluster protein depletion on SSU r-proteins could be explained by local assembly events involving direct protein-protein interactions.

**Fig 5.**
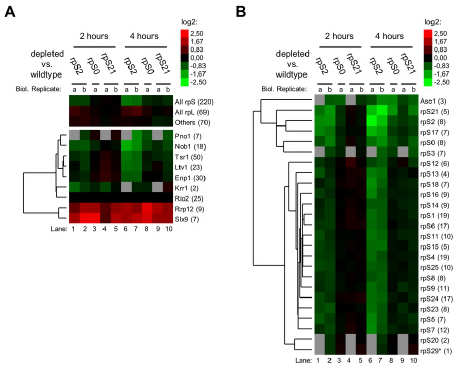
Semi-quantitative proteomic analyses of changes in the Rio2 associated SSU precursor protein composition after *in vivo* depletion of S0-cluster proteins. Yeast strains were cultivated for two and for four hours in glucose containing medium to shut down expression of either rpS0 (Y446), rpS2 (Y430), rpS21 (2704) or no r-protein (Y409). SSU precursors associated with TAP tagged Rio2 were affinity purified and the SSU precursor protein composition of each conditional expression mutant was compared with the one of wildtype cells (depleted vs. wildtype) by semi-quantitative mass spectrometry as described in Materials and Methods.Otherwise, see legend to Fig 4.

Proteomic analyses of the SSU biogenesis factor composition in Rio2 associated particles revealed several changes upon depletion of S0-cluster proteins (Fig 5A). Among all factors identified the binding of Nob1 and Pno1 to SSU precursors was consistently most severely affected. These proteins are direct neighbours of the S0-cluster in human late SSU precursors [54] and their proper integration into yeast SSU precursors might therefore be compromised upon failure of S0-cluster formation. In sharp contrast to the previous experiments a clear increase in levels of Rrp12 and of Slx9 was observed after expression shut down of any of the three S0-cluster proteins. Both of these ribosome biogenesis factors were previously suggested to facilitate the nuclear export of SSU precursors and to leave the cytoplasmic particles shortly afterwards by an unknown mechanism [11,12,16]. Accordingly, in agreement with the respective effects on Rio2 binding, the S0-cluster seemed to impact earlier steps of SSU maturation when compared to the S3-cluster.

### Impact of S0-cluster formation on the release of Slx9 from SSU precursors

To further corroborate these observations we genetically modified the conditional expression mutants of S0-cluster proteins and of rpS3 together with a corresponding wildtype strain to encode for TAP tagged Slx9. Affinity purifications using tagged Slx9 as bait were performed in mild buffer conditions and co-purifying (precursor-)rRNA was analyzed as before by northern blotting. As seen in Fig 6A in lanes 6-10 the apparent amounts of 20S pre-rRNA co-purifying with tagged Slx9 clearly increased upon *in vivo* depletion of S0-cluster proteins. In agreement with the previous proteomic analyses such an effect was not observed after expression shut down of rpS3. Hence, the results of these experiments indicated that the release of Slx9 is significantly and specifically delayed upon incomplete formation of the S0-cluster.

**Fig 6.**
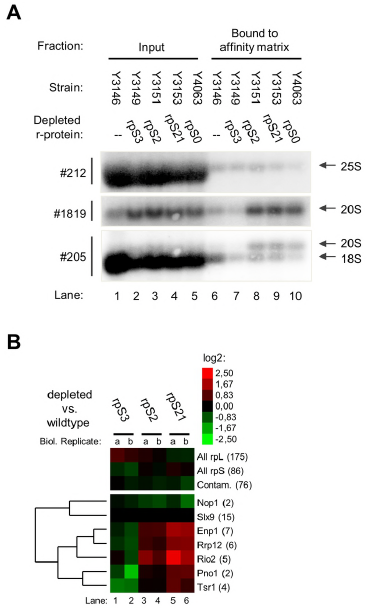
Association of Slx9 with SSU precursors depleted of S0- or S3-cluster proteins. In (A), the indicated yeast strains were cultivated for four hours in glucose containing medium to shut down expression of either rpS0, rpS2, rpS3, rpS21 or no r-protein. TAP tagged Slx9 was affinity purified from cellular extracts as described in Materials and Methods. Northern blotting analyses from input and affinity purified fractions are shown (see Materials and Methods). Oligonucleotides used for RNA detection are shown on the left (#, see S1 Fig for binding sites) and detected (pre-)rRNAs are indicated on the right. In (B), yeast strains were cultivated for four hours in glucose containing medium to shut down expression of either rpS2 (Y3151), rpS3 (Y3149), rpS21 (Y3153) or no r-protein (Y3146). SSU precursors associated with TAP tagged Slx9 were affinity purified and the SSU precursor protein composition of each conditional expression mutant was compared with the one of wildtype cells (depleted vs. wildtype) by semi-quantitative mass spectrometry as described in Materials and Methods. The results of two independent biological replicates (a, b) for ribosome biogenesis factors are visualized as heat map (colour legend in the upper right corner). Otherwise, see legend to Fig 4.

Pairwise comparison by semi-quantitative proteomics of Slx9 associated SSU precursors purified from wildtype strains with SSU precursors from mutant strains further confirmed this assumption (Fig 6B). Here, after *in vivo* depletion of the tested S0-cluster proteins rpS2 and rpS21 an increase in levels of factors associating with late SSU precursors as Rio2 and Enp1 and, to a lower degree of Pno1 and Tsr1, was observed. Likewise, the amount of Rrp12 was augmented as expected in case of a delay in its release. By contrast, a relative decrease in levels of all of these factors was observed in Slx9-TAP associated SSU precursors purified from cells in which expression of the S3-cluster protein rpS3 was shut down. Thus, these Slx9 interactome analyses were in agreement with an inefficient release of Slx9 and Rrp12 from SSU precursors upon incomplete assembly of the S0-protein cluster.

## Discussion

Dissociation of Rrp12 and Slx9 from precursor SSUs is thought to happen in wild type conditions shortly after nuclear export. The results of previous experiments analysing the subcellular steady state distribution of SSU pre-rRNA indicated that expression shut down of S0-cluster proteins leads to its accumulation in the cytoplasm [43]. Therefore, we think that the here observed delay in release of Slx9 and Rrp12 upon failure of S0 cluster formation likely occurs after nuclear export. Subcellular *in vivo* localisation of Rrp12 fused to green fluorescent protein in expression mutants of S0-cluster proteins were in agreement with this assumption (S2 Fig). In previous experiments addressing the dynamics of nuclear export a slowed kinetic was detected in S0-cluster mutants [43]. The lack of S0- cluster proteins on its own as well as the consequent impaired recycling of nuclear export factors Slx9 and Rrp12 or the inaccurate binding of Rio2 might account for this effect.

At this point there is no detailed structural information available on how yeast Slx9 and Rrp12 interact with SSU precursors. Still, recent cryo-EM studies revealed a partial structure of human Rrp12 bound to pre-SSUs [54]. Interestingly, human Rrp12 seems to keep the S0-cluster binding sites in the SSU head and body domain away from each other after the release of early nucleolar factors and the U3 sno-RNP. We therefore consider that the binding energy provided by stable association of the S0-cluster to all these sites might directly contribute to the release of Rrp12. Besides, evidence a role of Rio2 for the release of Rrp12 was previously observed in human cells [56]. If this release related role is conserved in yeast, the observed destabilised incorporation of Rio2 upon incomplete S0 cluster formation might compromise such a function. The pronounced role of the S0-cluster protein rpS2 in Rio2 binding might be related to its conserved beta-hairpin which closely approaches the Rio2 wHTH domain beta hairpin in SSU precursors [23]. In support of the functional relevance of these structural elements for SSU maturation, deletion of the yeast Rio2 wHTH domain is lethal and specific amino acid exchanges and deletions in the rpS2 beta hairpin do not interfere with rpS2 assembly but impair late SSU pre-rRNA processing [37,55,57].

In sum, the results of the herein described experiments helped to distinguish local clusters among the large group of late acting yeast SSU r-proteins which promote the stabilisation and processing of cytoplasmic SSU rRNA precursors in different ways. Formation of the S0-cluster seems important for the efficient release of two factors involved in nuclear export (Slx9 and Rrp12) as well as for the accurate incorporation of Rio2. In contrast, the results presented here corroborate the previously suggested role for S3-cluster proteins after Rio2 recruitment for the release of factors as Ltv1, Tsr1, Enp1 and Rio2 itself [31]. Besides, the r-protein assembly states detected in the respective conditional mutants differed, with major effects within each cluster, and some indication for a role of S0-cluster formation upstream of S3-cluster completion. The cluster-specific effects on binding and release of maturation factors indicate to us that, similar to what was observed for early nucleolar SSU maturation [52], several assembly checkpoints can be distinguished during late SSU maturation. Based on SSU precursor structure models we think that local assembly states can be sensed by specific biogenesis factors at these checkpoints. Possibly, direct factor interactions with the respective assembly sites are important for this, as discussed here regarding the S0-cluster for the Rio2 beta hairpin and for Rrp12 and as discussed in previous work regarding S3-cluster proteins and Ltv1. The respective maturation factors are either not efficiently released or recruited upon incomplete local assembly states and any downstream maturation step is thereby delayed. This might finally lead to incomplete protection of these SSU precursors from degradation and thereby promote selective accumulation of correctly assembled SSUs in the cytoplasm.

## Materials and Methods

### Yeast strains and microbiological procedures

Oligonucleotides, plasmids, and yeast strains used in this study are listed in S1-S3 Files. Genome editing of yeast strains for gene deletion, selection marker exchange or expression of chromosomally encoded tagged versions of biogenesis factors was done using PCR amplified DNA fragments with tag encoding elements and marker genes as described in [58–61]. Yeast transformation and selection was done as described in [60,62,63], for random spore analyses standard protocols were followed using mineral oil for spore enrichment [64]. Haploid cells were selected by resistance to canavanine and red colony colour due to deletion of the ADE2 gene. Expression of tag-fusion proteins was confirmed in strains with the expected genotypes by western blotting analyses.

For the experiments shown in Figs 1-5 strains logarithmically growing at 30°C in YPAG (1% yeast extract, 2% bacto peptone, 2% galactose, 100mg/l adenine hemisulfate) were cultivated for the indicated times at 30°C in YPAD (1% yeast extract, 2% bacto peptone, 2% glucose, 100 mg/l adenine hemisulfate).

### Semi-quantitative comparison of SSU precursor protein composition by mass spectrometry

Affinity purification of TAP tagged biogenesis factors with associated SSU precursors using rabbit immunoglobuline G coupled to magnetic beads and subsequent semi-quantitative mass spectrometric characterisation using iTRAQ™ 4plex reagents (Sciex) were performed as previously described [44]. The peak area for iTRAQ™ reporter ions were interpreted using the GPS-Explorer software package (Applied Biosystems) and Excel (Microsoft). The average iTRAQ™ ratio of all identified peptides (confidence interval > 95%) of a given protein was calculated (for r-proteins with two paralogues all peptides were combined), except for rpS21 and rpS29. Here, in experiments involving the respective conditional expression mutants the peptides specific to one of the two paralogues of each of these proteins were excluded from the analyses. All values obtained were normalized in regard to changes in the detected amounts of the bait protein used for affinity purification. Similarities in changes in levels of known ribosome biogenesis factors or SSU r-proteins identified by more than one peptide were further evaluated by hierarchical clustering analysis of datasets derived from several experiments. For this, proteins identified in more than 60% of the respective experiments were included and the cluster 3.0 software package was used with the city block distance and centroid linkage methods [65]. Data were visualized using Java Treeview (see http://www.eisenlab.org/eisen/?page_id=42).

### Affinity purification of TAP tagged ribosome biogenesis factors for subsequent RNA analyses

Affinity purification of tagged ribosome biogenesis factors on Immunoglobulin G sepharose and subsequent RNA analyses were performed as described in [44] with minor modifications. Cells from 500 milliliters of culture treated as described above were harvested by centrifugation for 6 minutes at room temperature at 5000g. The pelleted cells were washed once in ice cold water and were then stored at −20°C. Cells were thawed on ice, washed once in 5 milliliters of buffer A200 (200 mM potassium chloride, 20mM Tris pH8, 5mM magnesium acetate, 1 mM phenylmethylsulfonyl fluoride, 2 mM benzamidin) and were then suspended in 1.5 milliliters buffer A200 with 20U/milliliter RNasin (Promega) and 1mM dithiothreitol. 1.75g glass beads (0.75-1mm, Sigma) were added per milliliter of suspension and cells were disrupted by shaking them four times for 7 minutes at full speed on a Vibrax shaking platform (IKA) with cooling the samples in between on ice. The crude extract was cleared by two consecutive centrifugation steps for 5 and 10 minutes at 15000g at 4°C. The protein concentration of the resulting supernatant was determined by Bradford protein assay (Bio-Rad) and concentrations were adjusted with buffer A200 (with 20U/milliliter RNasin and 1mM dithiothreitol) to match among extracts of mutant and wildtype strains included in the respective analysis. Extract volumes containing 300 microgram of protein was added to 500 microliters cold buffer AE (50mM sodium acetate pH 5.3 and 10mM ethylenediaminetetraacetic acid) and stored at −20°C for subsequent RNA extraction and Northern blotting analyses. For each strain the same volume of extract was adjusted to 0.5% Triton X100 and 0.1% Tween 20 and added to 100 microliters of Immonuglobulin G sepharose 6 fast flow (Sigma-Aldrich) which was equilibrated in buffer A200+ (buffer A200 with 0.5% Triton X100 and 0.1% Tween 20). After incubation for one hour at 4°C the suspension was transferred to a poly-prep column (Bio-Rad) and washed two times with 2 milliliters A200+ followed by one washing step with 10 milliliters A200+. 80% of the suspension was transferred to a reaction tube and excess buffer was taken off after 2 minutes centrifugation at 2000g at 4°C. 500 microliters of cold buffer AE were added to the affinity matrix which was stored at −20°C until subsequent RNA extraction and northern blotting analyses.

### Sedimentation analyses on sucrose gradients

Sedimentation analyses on sucrose gradients were performed as described in [39] with minor modifications. Cellular extracts of yeast strains cultivated for four hours in YPAD were prepared in buffer A (20mM Hepes pH7.4, 5mM magnesium chloride, 100mM sodium chloride, 1 mM phenylmethylsulfonyl fluoride, 2 mM benzamidin, 3mM dithiothreitol, 10 Units per milliliter RNasin (Promega)) by agitation in the presence of glass beads as described above. Crude extracts were cleared by two consequent centrifugation steps at 4°C and 10000g for 5 minutes and 50 microliters of the resulting supernatant were loaded on a 10-50% (w/v) sucrose gradient which was prepared in buffer B (20mM Hepes pH 7.4, 5mM magnesium chloride, 100mM sodium chloride, 3mM dithiothreitol) using a gradient master device (Science Services). Centrifugation was done for 2 hours at 4°C and 40000 rounds per minute in a SW40 rotor (Beckman Coulter). Fractions of 500 microliters volume were collected. 50 microliters of each fraction were diluted 1:2 in buffer B, added to 500 microliters of cold buffer AE and stored at −20°C fur subsequent RNA analyses and northern blotting. For SDS PAGE and western blotting analyses 10 microliters of each fraction were diluted 1:2 in HU buffer (5% sodium dodecylsulfate, 200mM Tris pH 6.8, 1mM ethylenediaminetetraacetic acid, 1.5% (v/v) beta-mercaptoethanol, 8M urea, 0.002 % (w/v) bromophenolblue) and incubated at 65°C for 10 minutes.

### RNA extraction and northern blotting analyses

RNA extraction from samples taken during affinty purification was performed as described in [66] with minor modifications. Samples in buffer AE (see above) were thawed on ice and 500 microliters of phenol (equilibrated in buffer AE) and 50 microliters 10% SDS were added. After 6 minutes of rigorous shaking at 65°C the samples were cooled down on ice for two minutes and then centrifuged at 13000g and 4°C for two minutes. The upper layer was transferred to a new reaction tube containing 500 microliters of phenol (equilibrated in buffer AE) and after vortexing for 10 seconds the mixture was again centrifuged at 13000g and 4°C for two minutes. This procedure was repeated once with 500 microliters chloroform. The upper layer was again carefully taken off and RNA contained was precipitated by adding 2.5 volumes of ethanol and 1/10 volume of 3M sodium acetate pH5.3. In case of affinity purified fractions 10 microgram glycogen (Invitrogen) were added. Each sample was then briefly vortexed and incubated for more than 10 minutes at −20°C before centrifugation for 30 minutes at 4°C and 13000g. The supernatant was carefully discarded and the pellet was dissolved in 30 microliters of a buffer containing 6.5% formaldehyde, 50% formamide, 4mM sodium acetate, 10mM 3-(N-morpholino)propanesulfonic acid pH 7 and 0.5mM ethylenediaminetetraacetic acid) by shaking for 15 minutes at 65°C and subsequent cooling on ice. RNA was then size separated on 1.5% agarose gels in a buffer system containing formaldehyde and 3-(N-morpholino)propanesulfonic acid and was then transferred to positively charged nylon membranes (MP Biomedicals) as described in [67]. Hybridization with radioactively (^32^P) end labelled oligonucleotide probes indicated in the figure legends was performed overnight in a buffer containing 50% formamide, 0.75M sodium chloride, 75mM sodium citrate, 0.5% SDS, 1mg/ml Ficoll, 1mg/ml polyvinylpyrrolidone and 1mg/ml bovine serum albumine at 30°C. Washing was done for 15 minutes in 0.3M sodium chloride and 30mM sodium citrate and for 15 minutes in 0.15M sodium chloride and 15mM sodium citrate. Image plates (FujiFilm) were exposed to the radioactive signals on the washed membranes and were read out using phosphorimaging devices (FLA3000, FujiFilm, or Typhoon Imager FLA9500, GE Healthcare).

### SDS PAGE and western analyses

SDS PAGE and western transfer were performed according to standard protocols [67]. TAP-tagged proteins were detected after western transfer on membranes using a rabbit peroxidase anti-peroxidase (PAP) complex (Sigma-Aldrich) in a dilution of 1:5000. PAP was visualised by chemiluminescence and signals were detected by a fluorescence image reader (LAS3000, Fujifilm). Similarily, for testing the expression of fusion proteins with the eGFP_3xHA Tag the monoclonal 3F10 anti-HA antibody was used (Sigma-Aldrich) in a dilution of 1:1000 which was detected by 1:2500 diluted goat anti rat antibody coupled to peroxidase (Dianova).

### Life cell fluorescence microscopy

For life cell imaging strains logarithmically growing at 30°C in YPG (1% yeast extract, 2% bacto peptone, 2% galactose) were cultivated for four hours at 30°C in YPD (1% yeast extract, 2% bacto peptone, 2% glucose). Cells of 1 milliliter of culture were spun down in a micro centrifuge, re-suspended in 20 microliters of SDC with all supplements and directly observed with an Axiovert 200M fluorescence microscope equipped with a 63x Plan-Apochromat oil objective using filter set 46 (Zeiss).

## Acknowledgements

We thank former and present members of the chairs for biochemistry I and III for their constant support and for helpful discussions.

## Supporting Information

**S1 Fig. SSU pre-rRNA processing sites and major steady state SSU pre-rRNA intermediates in S. cerevisiae.** In (A) the primary transcript of 35S rRNA genes is schematically represented with 18S, 5.8S and 25S rRNA regions shown as grey boxes (not drawn to scale) and with external and internal spacers shown as continuous thin black line. Positions of major processing sites are highlighted by arrows and binding sites of oligonucleotides used in this study are indicated by black bars. In (B) the major steady state SSU pre-rRNA populations detected in S. cerevisiae are shown which result from co- or posttranscriptional processing events.

**S2 Fig. *In vivo* localization of Rrp12 after depletion of S0-cluster proteins.** The indicated yeast strains were cultivated for four hours in glucose containing medium (YPD) to shut down expression of either rpS0, rpS2, rpS21 or no r-protein. Cellular localization of GFP tagged Rrp12 as visualized by fluorescence microscopy (see Materials and Methods) is shown in (A) - (D). In (A) concentrated crescent shaped fluorescence signal, which is characteristic for the yeast nucleolus, is highlighted for two cells by yellow arrows.

S1 File. Oligonucleotides used in this study.

S2 File. Plasmids used in this study.

S3 File. Yeast strains used in this study.

S4 File. Mass spectrometry raw data and processed data for heatmaps

